# *Dnajc3* (HSP40) Enhances Axon Regeneration in the Mouse Optic Nerve

**DOI:** 10.1101/2024.10.08.617251

**Authors:** Jiaxing Wang, Ying Li, Felix L. Struebing, Sandra Jardines, Su-Ting Lin, Fangyu Lin, Eldon E. Geisert

## Abstract

A forward genetics approach was used to identify genomic elements enhancing axon regeneration in the BXD recombinant mouse strains. Axon regeneration was induced by knocking down *Pten* in retinal ganglion cells (RGCs) using adeno-associated virus (AAV) to deliver an shRNA followed by an intravitreal injection of Zymosan with CPT-cAMP that produced a mild inflammatory response. RGC axons were damaged by optic nerve crush (ONC). Following a 12-day survival period, regenerating axons were labeled by intravitreal injection of Cholera Toxin B (CTB) conjugated with Alexa Fluor 647. Two days later, labeled axons within the optic nerve were examined to determine the number of regenerating axons and the distance they traveled down the optic nerve. The analysis revealed a surprising difference in the amount of axonal regeneration across all 33 BXD strains. There was a 7.5-fold difference in the number of regenerating axons and a 4-fold difference in distance traveled by regenerating axons. These data were used to generate an integral map defining genomic loci modulating the enhanced axonal regeneration. A quantitative trait locus modulating axon regeneration was identified on Chromosome 14 (115 to 119 Mb). Within this locus were 16 annotated genes. Subsequent testing revealed that one candidate gene, *Dnajc3*, modulates axonal regeneration. *Dnajc3* encodes Heat Shock Protein 40 (HSP40), which is a molecular chaperone. Knocking down *Dnajc3* in the high regenerative strain (BXD90) led to a decreased regeneration response, while overexpression of *Dnajc3* in a low regenerative strain (BXD34) resulted in an increased regeneration response. These findings suggest that *Dnajc3* not only increases the number of regenerating axons, it also increases the distance those axons travel. This may prove to be critical for functional recovery in large mammals, where the distance axons travel to their target is considerably longer than that of the mouse. Thus, *Dnajc3* may play a critical role for functional recovery in humans by increasing the number of regenerating axons and the distance the regenerating axons travel.

## INTRODUCTION

Restoration of vision following injury to the optic nerve was once thought to be impossible, but is quickly becoming a reality. Under normal circumstances, injury to adult mammalian optic nerve results in axonal degeneration, retinal ganglion cell (RGC) death and a loss of function. The absence of regeneration is linked to epigenetics of transcriptional regulation[1–6]. Selected molecular process can be altered by experimental interventions designed to promote RGC survival and axons regrowth[3, 7–10]. Underpinning the RGC survival and axon regeneration is a cascade of cellular interactions involving multiple molecular pathways [3, 7, 10–20]. Ultimately, these pathways contribute to the survival of the injured RGC by modulating processes such as apoptosis[21, 22] and autophagy[7] and altering response to growth factors[16, 23–25]. Regenerating axons in the optic nerve not only requires the reactivation of axon growth programs[26], but also interact with growth inhibitors in the adult central nervous system (CNS). Inhibitory interactions are associated with many of the glial components within the optic nerve, such as astrocytes[27, 28], oligodendrocytes [15], and the glial scar[27, 28]. Ultimately, the regeneration is promoted by changes in transcriptional regulation[1–6] and signaling cascades[7, 29–31]. Given the complex nature associated with axonal regeneration, it is difficult to define the combination of events that would enhance the current protocols inducing axonal regrowth in the optic nerve.

One approach to facilitating axonal regrowth is to identify differences across inbred mouse strains or genetic backgrounds that enhance regeneration[13]. Omura et al.[13] tested 9 different inbred strains and found that axons of the CAST/Ei strain grew further in tissue culture on inhibitory substrates than the other 8 strains. In the intact mouse, the regenerative response of CAST/Ei strain was greater than that observed in the C57BL/6 strain. Unfortunately, using individual inbred mouse strains makes identifying the specific genomic loci driving increased regeneration very difficult. To overcome this problem, our group has taken advantage of the BXD genetic reference panel to identify genomic loci modulating induced optic nerve regeneration[32].

In the current study, a forward-genetics approach was used to define genomic loci modulating the axonal regeneration produced by knocking down *Pten* (phosphatase and tensin homolog)[7, 8] and inducing a mild inflammatory response[33] in mice that received a crush to the optic nerve. The amount of axon regeneration was quantified in 33 BXD recombinant inbred mouse strains. Defining the amount of induced axonal regeneration allows us to map a quantitative trait locus (QTL) modulating the regrowth of axons down the optic nerve[32]. This systems biology approach led to the identification of a specific gene within the QTL that enhances axonal regeneration, providing new insights into the genetic regulation of neuronal repair.

## MATERIALS AND METHODS

### Mice

For the mapping of a locus modulating axonal regeneration, 33 BXD recombinant inbred strains and their parental strains – C57BL/6J and DBA/2J were used in this study. For each strain, a minimum of 4 mice were used. All mice were 60-70 days of age at the time of initial treatment (See Supplemental Table 1). Controls were run with the C57BL/6J (n=6) and DBA/2J (n= 6) mice strains. For testing of candidate genes, the BXD strain with high regenerative responses we chose was BXD90 (n=21), ordered from The Jackson Laboratory. We used this strain to check the regeneration after modulating candidate genes. For testing of the low regenerating mouse strains, we ordered BXD34 mice (n=17) from The Jackson Laboratory. We also examined one parental strain C57BL/6J (n=20), ordered from The Jackson Laboratory. The mice were housed in a pathogen-free facility at Emory University, maintained on a 12-hour light/dark cycle, and provided with food and water ad libitum. All procedures involving animals were approved by the Animal Care and Use Committee of Emory University and were in accordance with the ARVO Statement for the Use of Animals in Ophthalmic and Vision Research.

### Surgery

The optic nerve regeneration protocol developed by others[7–9] was used to induce regeneration after optic nerve crush (ONC). The detailed protocol was described in our previous publication[32]. Briefly, the treatment included knocking down of *Pten* and intravitreal injection of Zymosan plus CPT-cAMP. We used AAV-sh*Pten*-GFP (*Pten* short hairpin RNA-GFP packaged into AAV2 backbone constructs, titer = 1.5×10^12^ vg/ml) to knock down *Pten*. The shRNA target sequence is 5’-AGGTGAAGATATATTCCTCCAA-3’ as described by Zukor et al.[34]. The efficient suppression of *Pten* expression in the retinal ganglion cells by this *Pten* shRNA has been proved in a previous study[32]. For all surgeries, the mice were deeply anesthetized with a mixture of 15 mg/kg of xylazine and 100 mg/kg of ketamine. Two weeks prior to ONC, the mice were deeply anesthetized and injected intravitreally with 2µL of AAV-sh*Pten*-GFP. Optic nerve crush was performed as described by Templeton and Geisert[35]. Briefly, under the binocular operating scope, a small incision was made in the conjunctiva. The optic nerve was visualized and then crushed 1 mm behind the eye with Dumont N7 angled crossover tweezers for 5 seconds, avoiding injury to the ophthalmic artery. Immediately following ONC, Zymosan (Sigma, Z4250, Lot#BCBQ8437V) along with the cAMP analog CPT-cAMP (Sigma, C3912, Lot#SLBH5204V, total volume 2µL) were injected into the vitreous to induce an inflammatory response and augment regeneration. The animals were allowed to recover from anesthesia and returned to their cages. Twelve days after ONC (two days before sacrifice), the animals were deeply anesthetized and Alexa Fluor® 647-conjugated Cholera Toxin B (CTB, ThermoFisher, Cat.#C34778) was injected into the vitreous for retrograde labeling of the regenerated axons. All the intravitreal injections and optic nerve crushes were performed by one well-trained postdoctoral fellow to avoid technical variation during the surgical procedure. At 14 days after ONC, the mice were deeply anesthetized and perfused through the heart with phosphate buffered saline (pH 7.3) followed by 4% paraformaldehyde in phosphate buffer (pH 7.3).

### Preparation of the optic nerve

Optic nerves along with optic chiasms and brains were dissected and post fixed with 4% paraformaldehyde in phosphate buffer overnight. The optic nerve was cleared with FocusClear™ (CelExplorer, Hsinchu, Taiwan) for up to 4 hours until totally transparent. A small chamber was built on the slide to provide enough space for the whole nerve thickness and to keep the nerve from being damaged from flattening. The optic nerve was then mounted in the chamber using MountClear™ (CelExplorer, Hsinchu, Taiwan) and the slides were cover-slipped. FocusClear™ has been used to clear brain tissue for whole brain imaging[36] as well as clearing of the optic nerve of transgenic zebrafish to observe axon regeneration[37]. It allowed us to scan the whole thickness of the optic nerve for better understanding of the status of axon regeneration, while providing clear imaging of regenerated axons from the optical slices scanned via confocal microscopy for counting. It also allowed us to determine the longest 5 axons or longest single axon along the nerve from z-stack of the whole nerve.

### Quantitation of axon regeneration

Cleared optic nerves were examined on a confocal microscope by scanning through individual optical slices. Green pseudo-color was used for CTB-labeled axons in all the optic nerve images of this study for clear visual observation. Stacked images were taken at 10µm increments, a total of 20-50 optical slices for each optic nerve. For quantifying the number of axons, we calculated the virtual thickness of an optical slice from the confocal microscope. As previously described[32], we determined that the thickness of the optical section was 6µm where refractive index (n) was 1.517, the numerical aperture (Na) was 0.45 and the excitation wavelength was 637nm. Since the optical section was 6µm and the spacing between optical sections was 10µm, single axons were not counted multiple times.

The number of CTB labeled axons at 0.5 mm from the crush site were counted in at least 6 sections per case and calculated by the equation (Σad=πr^2^ ∗ [average axons/mm]/t) as described by Leon et al. in 2000[33]. The cross-sectional width of the nerve was measured at the point at which the counts were taken and was used to calculate the number of axons per millimeter of nerve width. The total number of axons extending distance *d* in a nerve having a radius of *r*, was estimated by the summation of all sections. Since we used confocal images instead of longitudinal cross sections described in previous studies[8, 9], the optical resolution in *z* (0.5µm) was considered as the *t* (thickness of the slide) in the equation. The number of axons at 0.5mm from the crush site, the number of axons at 1mm from the crush site, the distance traveled by the 5 longest axons and the length of the longest single axon were all measured (See Figure 1 of our previous publication[32]).

**Figure 1.**
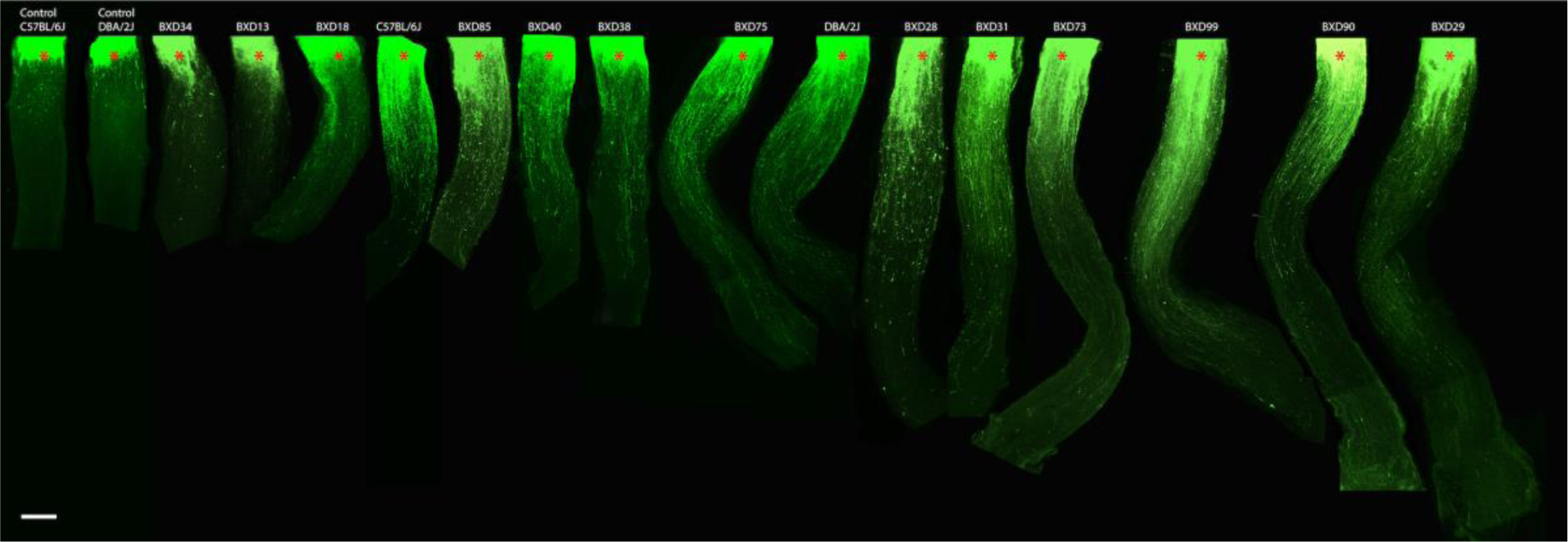
The figure is a series of photomicrographs from 17 optic nerves selected from 15 different strains of mice (N ≧ 4 in each strain). The two images to the left are from control mice with optic nerve crush only (Control C57BL/6J and Control DBA/2J). The remaining nerves are from animals in which the optic nerve was crushed and regeneration treatments were applied. Red asterisks represent the crush site. The strains from left to right are: C57BL/6J crush only, DBA/2J crush only, BXD34, BXD13, BXD18, C57BL/6J, BXD85, BXD40, BXD38, BXD75, DBA/2J, BXD28, BXD31, BXD73, BXD99, BXD90 and BXD29. The scale to the left represents 200µm.

### Interval Mapping

The axon regeneration data was subjected to conventional QTL analysis using simple and composite interval mapping using the mm10 assembly. Genotypes were regressed against each trait using the Haley-Knott equations implemented in the WebQTL module of GeneNetwork[38, 39]. Empirical significance thresholds of linkage were determined by permutations [40]. We generated interval maps for all 4 regeneration measures: number of axons at 0.5mm from the crush site, number of axons at 1mm from the crush site, distance traveled by the 5 longest axons and the length of the longest single axon. In the present paper, we made a synthetic trait for axon regeneration by combining all 4 datasets into a single measure and used this to produce an interval map. To identify loci, and also to nominate candidate genes, we used the following approaches: interval mapping for the traditional phenotypes, candidate gene selection within the QTL region, cis-eQTL analysis of gene expression, and trans-eQTL analysis.

### AAV vector

To knock down the genes of interest, we used a similar AAV vector to the *Pten* knock down as previously described[32]. Briefly, short hairpin RNA was designed for genes of interest, including *Dnajc3* (NM_008929.3) and *Uggt2* (NM_001081252.2), using the shRNA-designer from Biosettia (https://biosettia.com/support/shrna-designer/). The targeting sequence was selected on the exon region that contained no single nucleotide polymorphisms (SNPs) between C57BL/6J and DBA/2J strains. The shRNA sequence for the gene of interest was inserted into the AAV-sh*Pten*-GFP replacing the shRNA sequence for *Pten*. The transfection efficiency for these vectors is the same as AAV-sh*Pten*-GFP, which was tested previously[32]. For AAV-sh*Dnajc3*-GFP, the shRNA target sequence is 5’-GCAACCAGCAAATATGAAT-3’, virus titer = 1.6×10^14^ vg/ml. For AAV-sh*Uggt2*-GFP, the shRNA target sequence is 5’-GCCTGGGATTATCAGCAAT-3’, virus titer = 1.4×10^14^ vg/ml. For the overexpression of *Dnajc3* in the RGCs, the mRNA sequence of *Dnajc3* (NM_008929.4) was packed into the AAV vector. The virus titer was 4×10^12^ genomes. All of our plasmids were made at Emory Integrated Genomic Core (https://cores.emory.edu/eigc/) and were packed into AAV2 at Emory Viral Vector Core (https://neurology.emory.edu/ENNCF/viral_vector/). Previously[32], we have demonstrated that the AAV transduces approximately 54% of the retinal ganglion cells in mice.

### Testing Candidate Genes by Knockdown or Overexpression

Candidate genes were tested in our model optic nerve regeneration system. The AAV vector either with shRNA for knocking down a gene or with full-length sequence for overexpressing a gene were injected into the vitreous chamber one week before the start of the regeneration protocol. The vectors included: AAV-sh*Uggt2*-GFP and AAV-sh*Dnajc3*-GFP for knock down, or AAV-*Dnajc*3 for overexpression. For testing the effects of knocking down candidate genes, we used a BXD strain that demonstrated robust regeneration, BXD90. The BXD90 mice received an intravitreal injection of either AAV-sh*Uggt2*-GFP or AAV-sh*Dnajc3*-GFP. For testing the effects of overexpressing a gene, we used either a low regenerating strain, BXD34, or the parental strain, C57BL/6J. The AAV-*Dnajc3* vector was injected into the intravitreal chamber one week before the initiation of the regeneration protocol. The same strain of mice receiving AAV-GFP were used as a control group for both the knockdown and overexpression experiments.

### Statistical Analysis

Data are presented as Mean ± SE (Standard Error of the Mean). Differences in axon counts and regeneration distance between two strains were analyzed by Exact Wilcoxon-Mann-Whitney Test Calculator[41] (https://ccb-compute2.cs.uni-saarland.de/wtest/?id=www/www-ccb/html/wtest,). A value of *p* < 0.05 was considered statistically significant.

## RESULTS

Axon regeneration was examined in 33 strains of mice (31 BXD strains and the two parental strains, C57BL/6J and DBA/2J). Of these 33 strains, the data from nine were used in a previous study demonstrating that optic nerve regeneration is a complex trait[42]. The BXD strain set has proven to be a valuable genetic reference panel in vision research[43]. Here we examine the extent of axon regeneration 14 days following optic nerve crush. There was a surprising difference in regenerative capacity that was dependent upon the specific BXD strain examined (Figure 1). Some strains (BXD13, BXD18, and BXD34) showed very little axon regeneration; while other strains (BXD99, BXD90, and BXD29) displayed a significant amount of axon regrowth down the optic nerve (Figure 1). In all cases, optic nerve crush followed by the axon regeneration treatment led to more axonal regrowth than observed in the two control cases (C57BL6J and DBA/2J) which received optic nerve crush only (Figure 1). The axon regeneration for each strain was quantified by counting the number of regenerating axons at 0.5mm (Figure 2A) and 1mm (Figure 2B) from the crush site (See Supplemental Table 1). The distance the axons had traveled down the optic nerve was also quantified by measuring the length of the longest 5 axons (Figure 2C) and the distance traveled by the longest single axon (Figure 2D). These data were compared to axon regeneration in control mice that did not receive the regeneration treatment (Figure 2). When examining the regenerative capacity of the different BXD strains, the number of regenerating axons at 0.5mm varied by 7.5-fold, with the lowest number in strain BXD34 being an average of 135.4 axons and the greatest number of regenerating axons in strain BXD90 being an average of 1030 axons (Figure 2A). A similar pattern is observed 1mm from the crush site, with the lowest average number of regenerating axons in BXD18 and the highest average number of regenerating axons in BXD90 (Figure 2B). When examining the distance axons have regenerated down the optic nerve, the general pattern is similar to that observed for the number of regenerating axons. The fold change in axon growth for the longest 5 regenerating axons was 4 with the shortest axons in strain BXD34 and the longest axons in BXD29 (Figure 2C). This is also the case for the longest single axon with the shortest strain being BXD34 and the longest being BXD29 (Figure 2D). The data suggest that the number of regenerating axons and the distance the axons travel down the optic nerve reflect a similar genetic underpinning for the strains with the fewest axons also have the shorter regenerating axons. It is also the case that the strains with the greatest number of axons also have axons that travel the greatest distance down the optic nerve.

**Figure 2.**
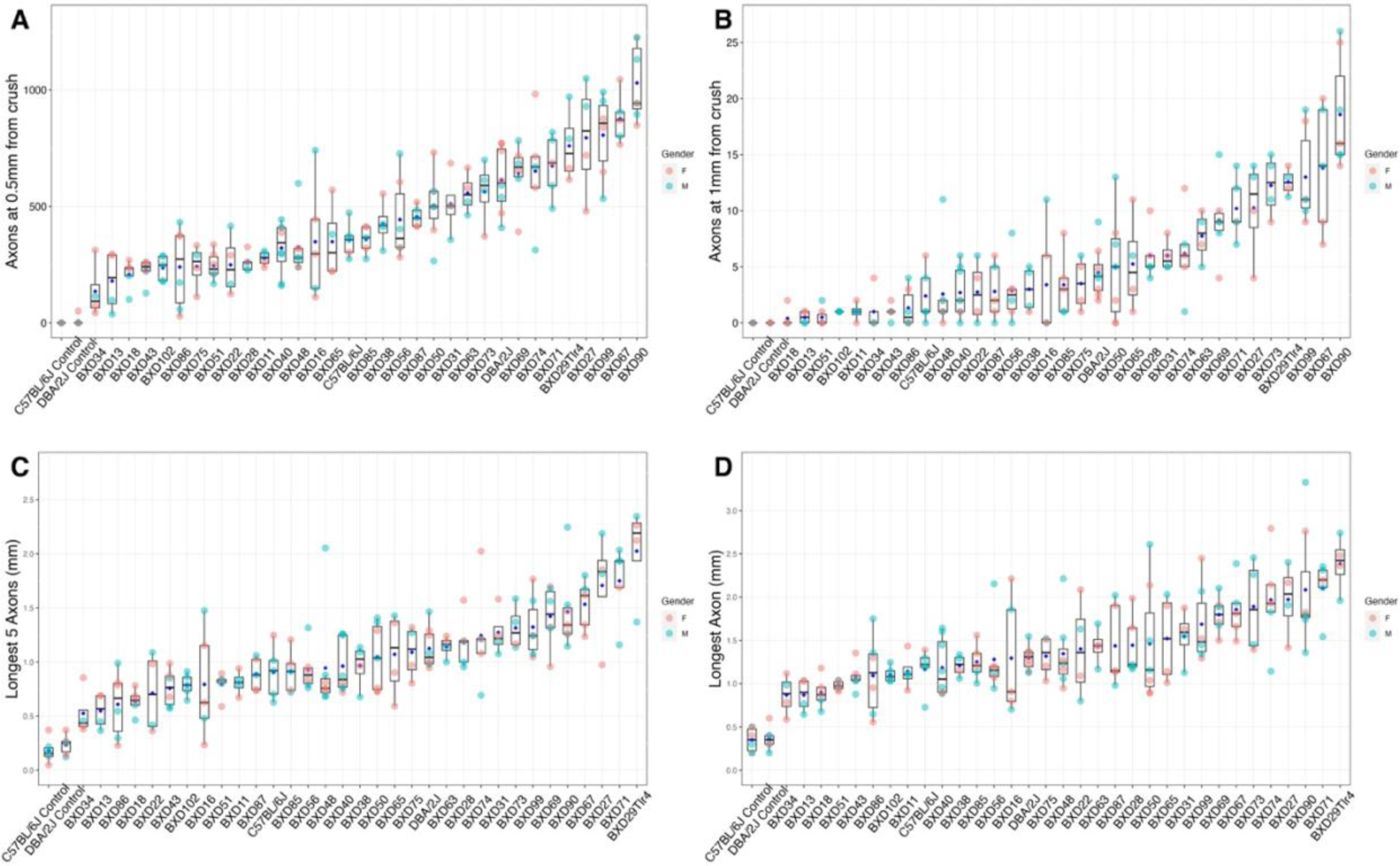
The number of axons at 0.5mm from the crush site (A), the number of axons 1mm from the crush site (B) are shown. In addition, the distance traveled by the longest 5 regenerating axons (C) and the longest single axon (D) is illustrated, in two control strains (C57BL/6J and DBA/2J untreated mice) and in 33 strains that went through the regeneration protocol. Each animal is represented by a single dot, with the pink dots for females and the light blue dots for males. Comparing the strains with the greatest regeneration with the strains with the least regeneration, there is a 7.5-fold difference in the number of axons at 0.5mm from crush site (A) and 38-fold difference at 1mm from crush site (B). There is an approximate 4-fold difference in the distance of the longest 5 axons and 3-fold difference in the distance of the longest single axon, when comparing the strains with longest axons to the ones with the shortest axons. The fact that the parental strains are not the extremes of the distributions indicates that there is genetic transgression in axonal regeneration, suggesting a minimum two genomic loci modulating the phenotype of axon regeneration.

### Interval Mapping

These data were used to map genomic loci that may modulate the regenerative response in the BXD strains. The unbiased, forward genetics approach was used to create a genome-wide interval map for each of the measures of regeneration: number of axons 0.5mm from the crush site, number of axons 1mm from the crush site, distance traveled by the 5 longest axons and the length of the longest single axon (Supplemental Figure 1). All four interval maps showed a similar pattern with a large peak being on distal Chromosome (Chr) 14. The peak on Chr 14 (115 Mb to 119 Mb) is above the suggestive level for all measures. The map on the longest single axon has a QTL that reached the significance level (Supplemental Figure 1D). Since all of the genome wide scans were similar, all of these data were combined to create a synthetic trait, “axonal regeneration”. The genome-wide scan for axonal regeneration revealed the same significant peak on Chr 14 (Figure 3). For further analysis, we used data from the combined synthetic trait, axon regeneration, to identify the QTL modulating axon regeneration as well as defining the candidate genes within the region.

**Figure 3.**
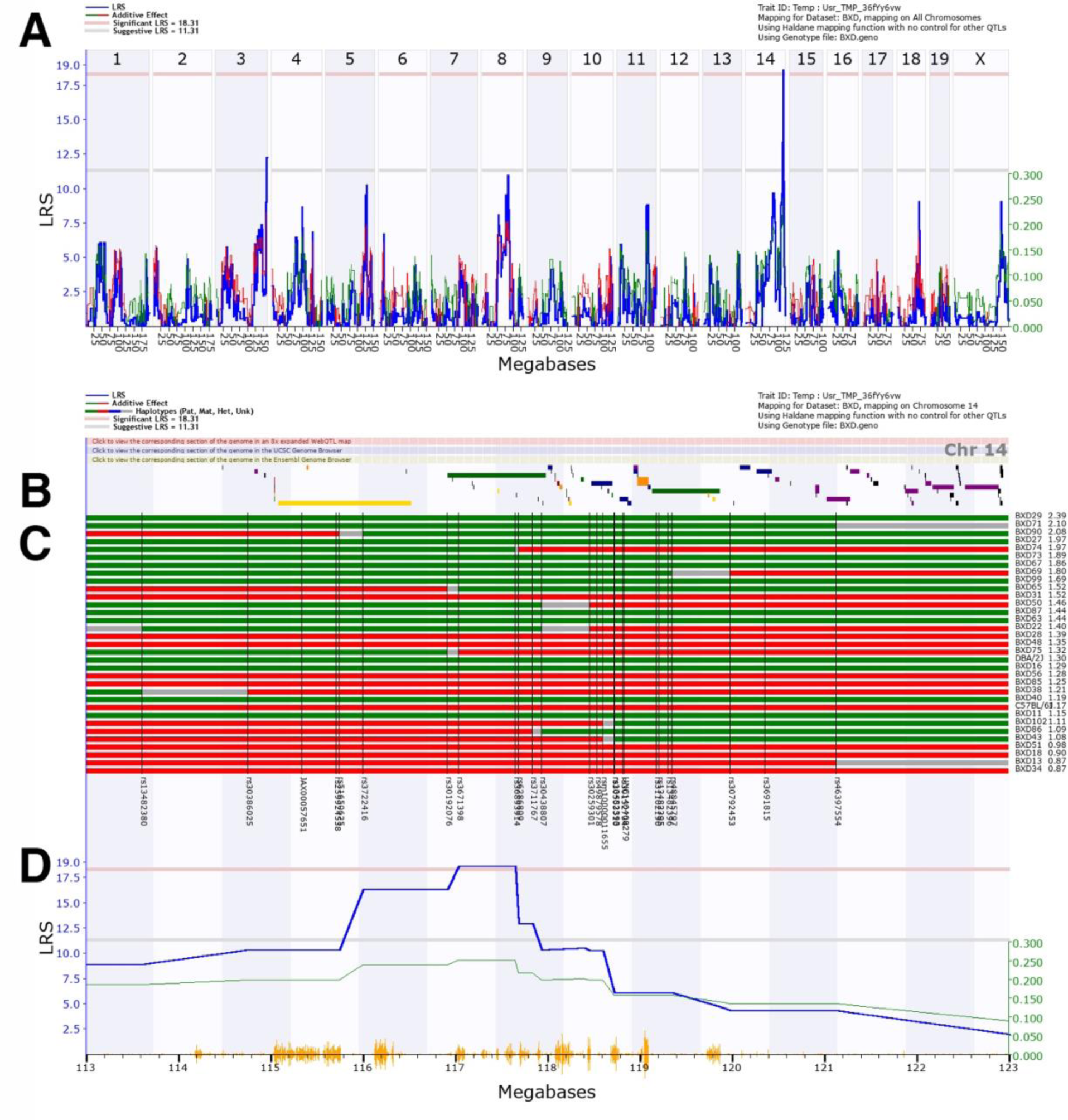
A genome-wide interval map of axon regeneration (A). The interval map plots the likelihood ratio statistic (LRS) across the genome from chromosome 1 to chromosome X. The light gray line is the suggestive level and the light red line is the genome-wide significance (*p* = 0.05). When the optic nerve regeneration measures were mapped to the mouse genome, there was a significant association between optic nerve regeneration and a locus on Chromosome 14. A map of gene locations across Chr 14 from 113 to 123Mb is shown (D). The locations of genes within this region are schematically shown in the top panel (B). The haplotype map for the 33 strains in the optic nerve regeneration dataset is illustrated in the middle panel with each strain indicated on the right (C). Red represents the B6 alleles, green defines the D2 alleles, blue represents regions of the DNA that are heterozygotic and gray is unmapped. The genomic markers used in the mapping process are listed at the bottom of the haplotype map. At the far right is a list of the specific BXD strains and the associated regeneration measurements going from the most regeneration at the top to the least at the bottom. A QTL map of regeneration on Chr 14 is shown below the haplotype map. An LRS above the pink line is statistically significant (p<0.05). A positive additive coefficient (green line) indicates that D2 alleles are associated with higher trait values. The significant peak for optic nerve regeneration is from 114 to 119 Mb. Vertical orange lines at the bottom of the plot show the SNPs on Chr 14.

### Candidate Genes

The interval map for the longest single axon and the synthetic trait axon regeneration both had a significant QTL peak on Chr 14. To identify genes modulating axonal regeneration in the BXD strains, we examined a 4 Mb region around the peak ranging from 115 Mb to 119 Mb (Figure 3). Within this region, there were 16 annotated genes: *Mir18*, *Mir19a*, *Mir20a*, *MIr17hg*, *Gpc5*, *Gpc6*, *Gm19845*, *Dct*, *Tgds*, *Gpr180*, *Sox21*, *Abcc4*, *Cldn10*, *Dzip1*, *Dnajc3* and *Uggt2*. In general, ideal candidate genes should have a nonsynonymous SNP between the two parental strains that affects protein sequence and function, or there should be genomic elements with cis-QTLs affecting expression levels. Of the 16 genes within the region, two had nonsynonymous SNP: *Abcc4* and *Cldn10*. To determine if any of these amino acid changes would affect protein function, we ran a SIFT analysis[44]. All of the changes in both genes were tolerated and should not disrupt protein function. Thus, these two genes did not appear to be good candidates and were removed as potential candidate genes. Two genes were found with cis-QTLs: *Dnajc3* and *Uggt2*. The difference in expression levels of these two candidate genes was confirmed by examining RNAseq comparisons between the parental strains[45]. Furthermore, both of these genes are expressed in most of the retinal ganglion cell types in single cell RNAseq datasets[46]. These data indicate that both *Dnajc3* and *Uggt2* are good candidate genes for modulating axonal regeneration.

### Knockdown of *Dnajc3* and *Uggt2*

To further investigate the roles of the two candidate genes in optic nerve regeneration, we extended our study using the existing regeneration model. As an initial test, we chose one of the high regenerative strains, BXD90, to examine the effects of knocking down gene expression using AAV delivery of shRNA. AAV-shRNA directed against either *Uggt2* or *Dnajc3* was injected into the vitreous chamber of the mouse eye, knocking down the expression of each candidate gene. The injection was given a week prior to *Pten* knockdown (Figure 4A). This allowed the effective transduction of cells with AAV carrying candidate gene-specific shRNA. This also produced a decrease in gene expression prior to the beginning of the experiment without disrupting the established neural regeneration experimental protocol. The control group received AAV-GFP at the same time point. A successful knockdown of the targeting gene *Dnajc3* is demonstrated in Figure 4 (B-D).

**Figure 4.**
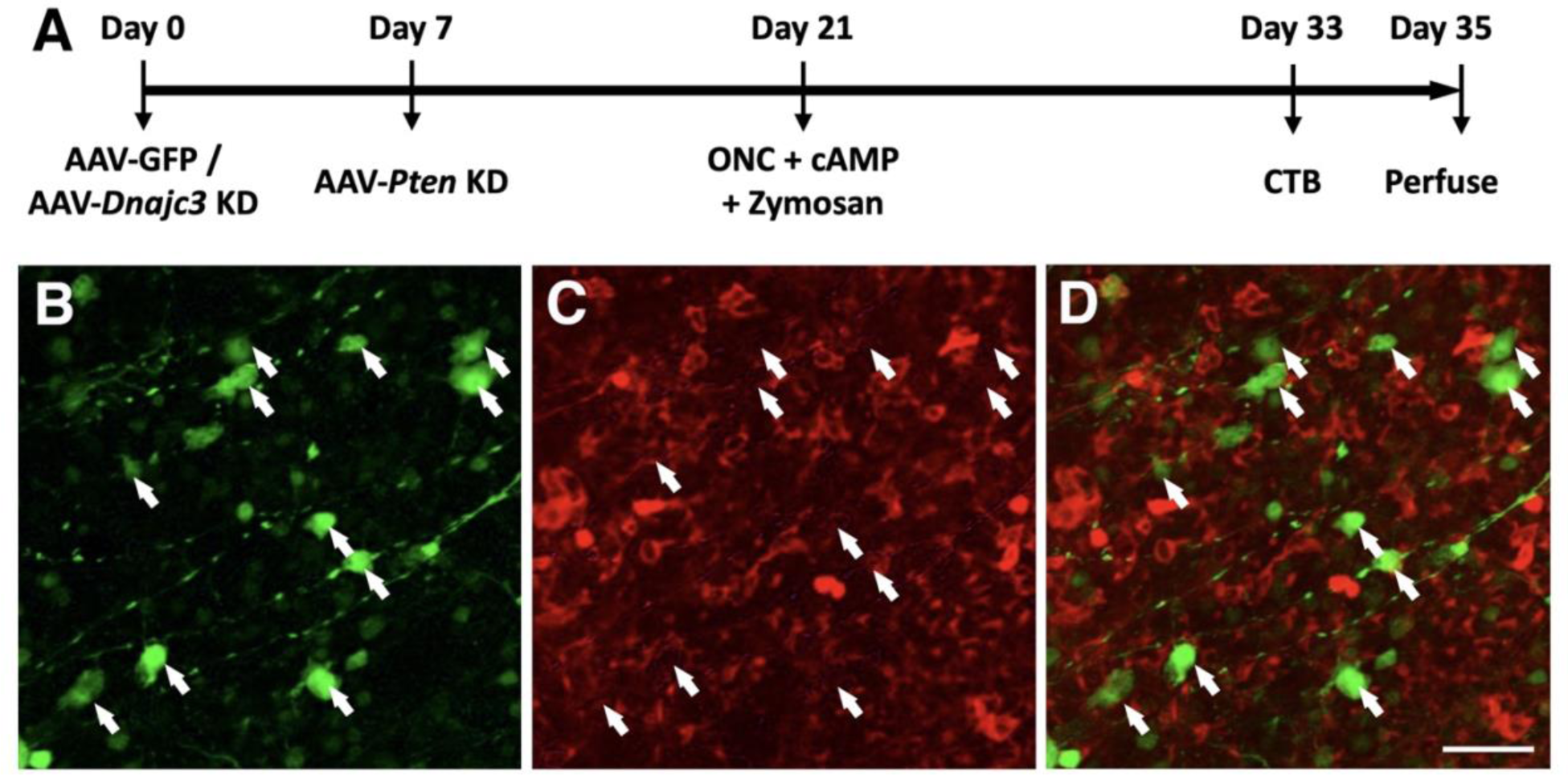
The modified regeneratin protocol is illustrated in A, 7 days prior to the knockdown of *Pten* (Day 0), AAV-*Dnajc3* KD or AAV-GFP (control) was injected into the eye. At day 7, AAV-sh*Pten*-GFP was then injected into the eye. Two weeks later (Day 21), the optic nerve is crushed and Zymosan and cAMP were injected into the eye. On day 33, the eye was injected with Alexa Fluor® 647-conjugated CTB and two days later the animal was perfused. In panels B-D, flatmount of a retina injected with AAV-sh*Dnajc3*-GFP are shown. In the first panel B, the staining for GFP is shown labeling RGCs. Panel C shows the labeling for HSP40 (*Dnajc3*). Notice the lack of HSP40 staining in the GFP expressing RGCs (white arrows in B-D). The final panel D is a merged image of panels B and C. The scale bar in D represents 50µm.

Our results showed that selectively knocking down *Uggt2* in RGCs did not impact optic nerve regeneration (Figure 5 A-D), making it unlikely to be the gene modulates axon regeneration in the BXD strains. Knocking down of *Dnajc3* in RGCs decreased the number of axons regenerating in the optic nerve (Figure 5A and 5B) as well as the distance that the axons traveled down the optic nerve (Figure 5C and 5D) after regeneration treatment for ONC. Quantifying the regenerative capacity revealed a significant 24.6% decrease (767.6±44.5 vs 1017.9±46.5, p < 0.05) in the number of axons at 0.5mm from the injury site compared with the GFP group. There was also a significant 53.7% decrease (8±1 vs 17.3±1.6, p < 0.05) at 1mm from the crush site. A similar result was observed in the distance axons regenerated down the optic nerve. There was a 28.5% decrease in the distance of the 5 longest axons in the *Dnajc3* knockdown mice relative to the GFP controls (1±0.1 vs 1.4±0.1, p < 0.05), and for the longest single axon there was a 38.5% decrease in distance traveled (1.3±0.1 vs 2±0.2, p < 0.05). Detailed data is shown in Supplementary table 2. These data indicate that *Dnajc3* is a good candidate gene for modulating axon regeneration down the optic nerve.

**Figure 5.**
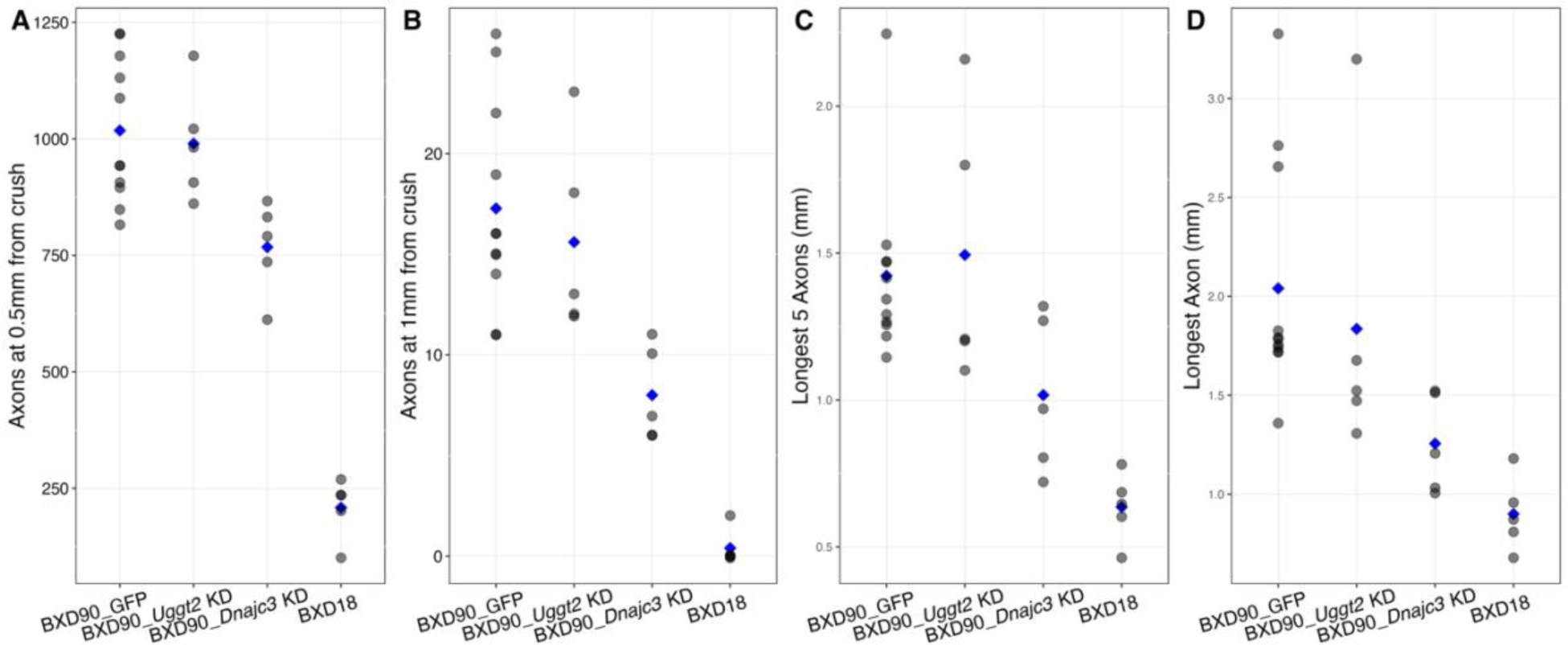
*Uggt2* and *Dnajc3* knockdown in BXD90. BXD90 was one of the strains with robust optic nerve regeneration induced by our regeneration protocol. We used AAV-shRNAs to knockdown the two candidate genes, *Uggt2* and *Dnajc3*, in RGCs prior to testing for optic nerve regeneration. Each animal is represented by a dot and the means are presented as blue diamonds. In panel A, the number of axons at 0.5mm from the crush site is shown. The first column is the data from control mice receiving AAV-GFP. The effects of knocking down *Uggt2* is shown in the next column (BXD90_*Uggt2*_KD). The data from knocking down *Dnajc3* is shown next (BXD90_*Dnajc3* KD). Notice a significant decrease in the number of axons regenerating (p < 0.05 Mann-Whitney U test). The data from one of the low regenerating strains (BXD18) is also shown for comparison. Similar results are shown for number of axons at 1mm from the crush site (B). This is also the case for the distance axons regrew down the optic nerve, for both the 5 longest axons (C) and the longest single axon (D). These data demonstrate that knocking down *Uggt2* does not affect axon growth in BXD90. It also demonstrates that knocking down *Dnajc3* can modulate the amount of axon regeneration. Thus, *Dnajc3* is the best candidate at the Chr 14 locus for modulating axonal regeneration.

The axonal regeneration observed following the knockdown of *Dnajc3* in BXD90, a high regenerative strain, is significantly greater than that observed in BXD18, a low regenerative strain (Figure 5). This is the case for the number of regenerating axons as well as the distance the axons travel down the optic nerve. Although the regenerative capacity is significantly diminished in *Dnajc3* knock down in BXD90 mice relative to the GFP controls (p < 0.05 Mann-Whitney U test), it still does not reach the level of the low regenerative strain, BXD18 (Figure 5). These findings indicate that *Dnajc3* modulates optic nerve regeneration in the BXD strain set; however, *Dnajc3* is not solely responsible for the variation in regeneration observed in the BXD strain set, for the increased regeneration is not completely blocked by knocking down *Dnajc3*.

### Overexpression of *Dnajc3*

To provide additional evidence that *Dnajc3* modulates axonal regeneration, we used AAV to overexpress *Dnajc3* in the RGCs of one of the parental strains (C57BL/6J) and one of the lowest regenerating strains (BXD34). The full cDNA sequence of *Dnajc3* was placed into a AAV vector and packaged into AAV2 capsid. The AAV-*Dnajc3* was injected into the vitreous of 10 C57BL/6J mice and 9 BXD34 mice, while the controls were injected with AAV-GFP in 10 C57BL/6J mice and in 8 BXD34 mice one week prior to the optic nerve regeneration protocol (similar schedule as shown in Figure 4A). In both C57BL/6J and BXD34 strains, there was a significant increase (p<0.05 Mann-Whitney U test) in the number of regenerating axons (Figure 6 A and B) and the distance the regenerating axons traveled down the nerve (Figure 6 C and D). Detailed data is shown in Supplementary table 2. Thus, overexpressing *Dnajc3* in low regenerating strains increased the amount of axon regeneration down the optic nerve. Taken together, the decrease in axonal regeneration following knocking down *Dnajc3* and the increase in regeneration following overexpressing *Dnajc3*, demonstrates that one of the genomic elements modulating axonal regeneration in the BXD strain set is *Dnajc3*.

**Figure 6.**
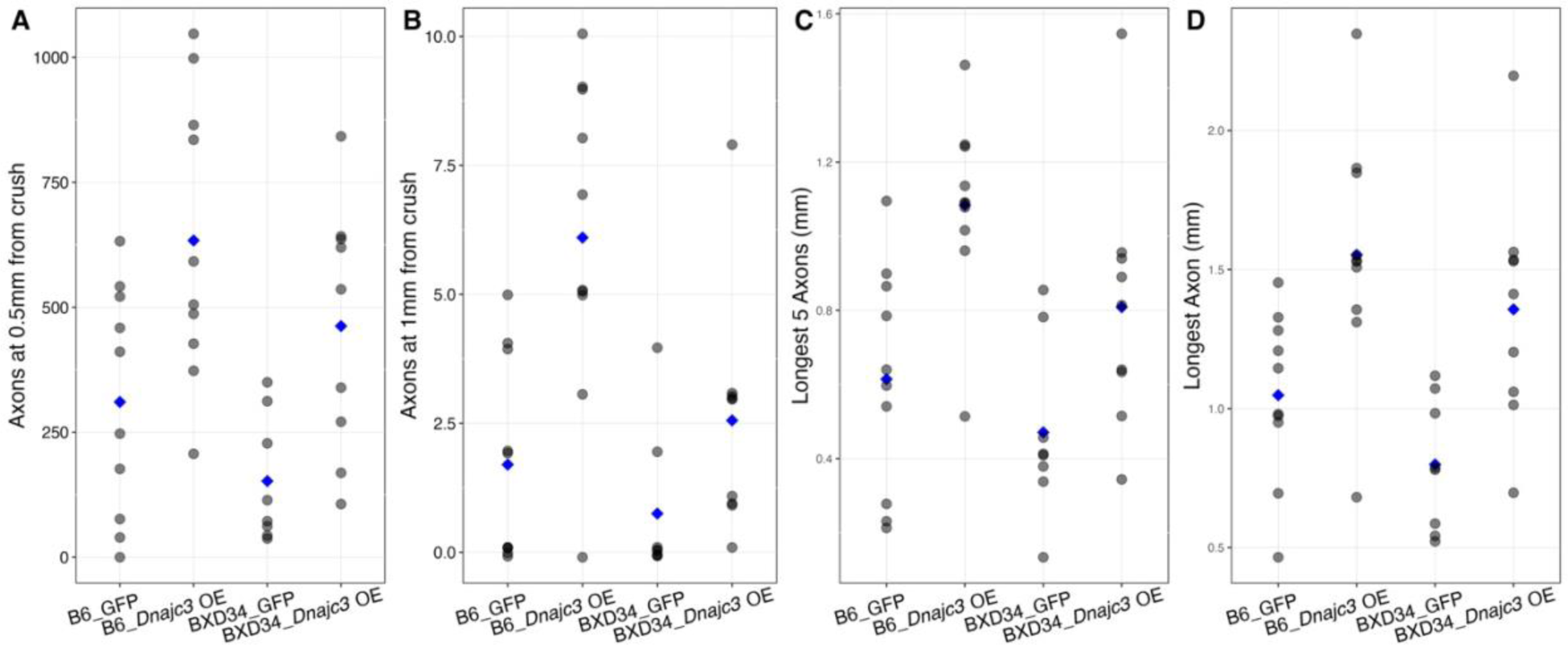
Overexpression of *Dnajc3* in C57BL/6J mice and in BXD34 mice. BXD34 was one of the strains with least optic nerve regeneration induced by our regeneration protocol. *Dnajc3* was over expressed using AAV-*Dnajc3* to express it in RGCs prior to testing for optic nerve regeneration. Each animal is represented by a dot and the means are presented as blue diamonds. In panel A, the number of axons at 0.5mm from the crush site is shown. The first column is the data from control C57BL/6J mice receiving AAV-GFP (B6_GFP). The effects of overexpressing *Dnajc3* are shown in the next column (B6_*Dnajc3*_OE). There was a significant (p < 0.05 Mann-Whitney U test) increase in the number of axons when *Dnajc3* was overexpressed. The next column shows the data from the BXD34 GFP control (BXD34_GFP). The data from overexpressing *Dnajc3* in BXD34 is the last column (BXD34_*Dnajc3*_OE). Notice a significant increase in the number of axons regenerating (p < 0.05 Mann-Whitney U test). Similar results are shown for number of axons at 1mm from the crush site (B). This is also the case for the distance axons regrew down the optic nerve, for both the 5 longest axons (C) and the longest single axon (D). These data demonstrate that overexpressing *Dnajc3* in a low regenerating strain of either C57BL/6J or BXD34 increases the regeneration of axons following optic nerve crush.

## DISCUSSION

In the present study, we have used a forward-genetics approach to identify a genomic locus that modulates the regenerative response induced by knocking down *Pten* and creating a mild inflammatory response with Zymosan and CPT-cAMP. This approach identified one locus on Chromosome 14 (115 Mb to 119 Mb) that modulates optic nerve regeneration. Within this locus, there were 16 annotated genes, two of which were good candidate genes (*Uggt2* and *Dnajc3*). Subsequent testing eliminated *Uggt2* as a modulator of axonal regeneration and demonstrated that changes in expression levels of *Dnajc3* can alter the regenerative response. Knocking down *Dnajc3* in a high regenerating strain of mice decreases axon regeneration; while, overexpressing *Dnajc3* in a low regenerating strain increases the ability of axons to regenerate in the optic nerve. In the BXD strain set, *Dnajc3* modulates the regenerative response produced by knocking down *Pten* and inducing a mild inflammatory response with Zymosan and CPT-cAMP.

*Dnajc3* is expressed in RGCs at relatively high levels and is localized to the endoplasmic reticulum (ER)[47]. *Dnajc3* encodes p58IPK, also known as heat shock protein 40 (HSP40)[48, 49]. Like many heat shock proteins, HSP40 is an ER chaperone playing a role in the unfolded protein response (UPR) when neurons are stressed[48, 49]. The UPR plays an essential role in maintaining cellular homeostasis in CNS neurons by activating in stress conditions when there’s an accumulation of misfolded or unfolded proteins in the ER. Unlike many of the chaperone proteins, HSP40 can also function as an inhibitor of eIF2a, a protein kinase localized to the inner surface of the endoplasmic reticulum[49]. Crucially, in the context of retinal health, the role of HSP40 (p58IPK) in protein homeostasis and anti-inflammatory response has been identified as protective against retinal ganglion cell degeneration[47, 50, 51], a primary cause of visual impairment in conditions like glaucoma. The protective effect of HSP40 was revealed by Zhang’s group from both in vitro[47, 51] and in vivo[50, 51] experiments. They reported that deficiency of HSP40 makes the RGCs more susceptible to cell death by the ER stress inducer; while overexpression of p58IPK by AAV increases RGC survival under ER stress[51]. HSP40 protects from retinal ischemia/reperfusion and from elevated intraocular pressure (IOP) caused by microbead injections into the anterior chamber[51]. Knocking down HSP40 in vivo and in vitro causes a decrease in RGC survival while overexpressing the protein promotes neuronal survival in culture[51]. These data strongly support the role of HSP40 in protecting RGCs from stress induced cell death.

Using the BXD mouse strains, we show that *Dnajc3* is a genomic element modulating axonal regeneration. This includes the number of regenerating axons at 0.5mm and 1mm from the crush site, as well as the distance these axons travel down the optic nerve, whether it is the 5 longest axons or the single longest axon. Our results suggest one potential mechanism responsible for the partial increase in the number of regenerating axons is the neuroprotection of RGC cell body from death. Previous studies[47, 50] demonstrate that overexpressing *Dnajc3* (HSP40) is neuroprotective for RGCs in an elevated IOP model of glaucoma and in an ischemia/reperfusion model in the mouse. If *Dnajc3* can act as a neuroprotective genomic element, then the increase in the number of regenerating axons could be directly due to the fact that more RGCs are surviving the insult to the optic nerve. Thus, the number of regenerating axons may be directly related to the neuroprotective effects of *Dnajc3* in RGCs; the more RGCs that survive following optic nerve crush, the more axons that are available to regenerate down the optic nerve.

Interestingly, we have found that the genomic loci modulating the distance axons regenerate down the optic nerve are similar to those affecting the number of regenerating axons. Clearly, RGC survival is one of the fundamental aspects necessary for axon regeneration, for if the cell body dies, the cell will not be able to regrow down the optic nerve. That being said, there is increasing evidence that RGC survival and axon regeneration can be two completely independent processes. An example of this is the effect of *Sox11* on retinal injury. The overexpression of *Sox11* in mouse RGCs promotes regeneration[10]. Interestingly, it does not promote regeneration in some of the RGC subtypes. αRGCs, which are normally associated with induced regeneration down the optic nerve, are killed by *Sox11* overexpression[10]. Thus, the increased axonal regeneration induced by *Sox11* overexpression must be due to a response from a different RGC subtype, not the αRGCs. The downregulation of *Sox11* increased RGC survival following injury of optic nerve axons[52, 53]. Even though there was an increase in RGC survival, there was not an improved regeneration of axons down the optic nerve in an induced optic regeneration model[52, 53]. Thus, the neuroprotection of RGCs does not result in an obligatory increase in axonal regeneration. That being said, there may be elements that facilitate RGC survival and facilitate axonal regeneration.

Independent of RGC survival, HSP40 (p58IPK) might also influence axon regeneration in the following aspects. In the past, it was believed that while neurons in the peripheral nervous system (PNS) could regenerate after injury, mature neurons in the CNS could not. However, numerous subsequent studies have achieved regeneration of CNS neurons through experimental methods. In the injured PNS, regeneration involves the local synthesis of damage signals and translation of growth-promoting mRNA, such as Gap-43 and β-actin[54]. Under certain conditions, similar mRNA was also detected in injured CNS axons[55]. Local protein synthesis events detected in embryonic axons and adult PNS axons play a pivotal role in neuronal regeneration[56]. For CNS neurons, axon regeneration requires the fulfillment of several conditions. One critical aspect is the synthesis and transportation of proteins to supply the nutrients and raw materials necessary for axon regeneration. First, protein synthesis is vital for the formation of growth cones. Research has shown that inhibiting protein synthesis impairs growth cones, which are essential structures for axon regeneration[57]. Second, processes involved in axon regeneration, such as retrograde signaling, growth cone formation, and axon elongation, all depend on local translation[56]. Adult local transcriptomes detected in retinal ganglion cells, as well as translation mechanisms in several adult brain regions, have confirmed local translation in mature CNS axons[58–60]. Moreover, during CNS development, ribosome localization in axons might gradually decrease, indicating that ribosome positioning and mRNA translation specificity change with maturity[58, 61]. After injury, the local translation dynamics of adult CNS axons might also decline, such as reduced axonal transcripts or a lack of functional ribosomes[62]. Furthermore, local translation might be hindered due to incorrect mRNA transport or the retention of mRNA in stress granules[54, 63]. Therefore, it can be inferred that the limited regenerative capacity of CNS axons might be related to reduced local translation. Supporting this, earlier research by Park and colleagues[7] demonstrated that enhancing protein synthesis by augmenting mTOR signaling activity through silencing the *Pten* gene greatly promotes optic nerve axon regeneration, possibly by influencing local axonal translation[7]. In this experiment, the establishment of the axon regeneration model was based on silencing the *Pten* gene. As a chaperone protein in ER stress, HSP40 may alleviate ER stress, thereby enhancing the protein synthesis capability of the ER as well as the protein translation and transportation abilities at the site of axon injury. This series of effects means that neurons influenced by HSP40 not only have enhanced survival capabilities but also exhibit significantly increased axonal regenerative abilities.

### *Dnajc3* is not the only modulator of axon regeneration in the BXD strains

When we examine the effects of altering the expression of *Dnajc3*, it is clear that it is not the sole element affecting the difference in axonal regeneration across the BXD strain examined in the current study. Knocking down *Dnajc3* in a strain with extensive axonal regeneration does not reduce the axonal regrowth to the level of the lower expressing strains. Furthermore, overexpressing *Dnajc3* in a strain with limited axonal regrowth does not enhance the regeneration to the same level as a strain with robust optic nerve regeneration. Taken together, these data indicate that there is at least a second genomic element interacting with *Dnajc3* or directly modulating optic nerve regeneration that differs between the two strains of mice.

## CONCLUSIONS

Optic nerve regeneration is a highly complex process with many factors interacting and influencing each other. Using a forward-genetics approach, we identified *Dnajc3* as a genomic element promoting an increase in the number of regenerating axons as well as the distance these axons travel. The protein it encodes, HSP40, is one element modulating axonal regeneration in the mouse. The gene product of HSP40 is an important factor within the retinal ganglion cells, and the protein either overexpressing or knocking down its expression affects the regenerative capacity of injured axons in an experimentally induced optic nerve regeneration model. One of the underlying mechanisms in this increased regeneration is probably due to its role in protecting RGCs from death. In addition, it may improve the ability of neurons to overcome the unfolded protein response that complicates axons regrowth. Together, the actions of HSP40 may be critical to functional regeneration in humans as the distance the axons have to travel to reach their targets in the human brain is considerably longer than that in most rodent models.

## List of Abbreviations

AAV: Adeno Associated Virus
ONC: Optic Nerve Crush
RGC: Retinal Ganglion Cell
GFP: Green Fluorescent Protein
HSP40: Heat Shock Protein 40
QTL: Quantitative Trait Locus
CTB: Cholera Toxin B
Chr: Chromosome
UPR: Unfolded Protein Response
CNS: Central Nervous System
PNS: Peripheral Nervous System
ER: Endoplasmic Reticulum
IOP: Intraocular Pressure
LRS: Likelihood Ratio Statistic
SNP: Single Nucleotide Polymorphism

## Acknowledgements

The authors would like to thank Rebecca King for her technical assistance in this study. We thank the Emory Viral Vector Core for the production of AAV (NINDS Core Facilities Grant P30NS055077) and the Emory Integrated Genomics Core (subsidized by the Emory University School of Medicine and NIH UL1TR002378). The content is solely the responsibility of the authors and does not necessarily reflect the official views of the National Institutes of Health.

## Authors Contributions

The following are the authors contribution to the publication: design of experiments (JW, YL, FLS, EEG), provide funding (JW, EEG), collection and analysis of data (JW, YL, FLS, SJ, S-TL, FL, EEG), writing the manuscript (JW, EEG), and making substantial edits to the manuscript (JW, S-TL, FL, EEG).

## Funding

This study was supported by two grants from the BrightFocus Foundation G2019111 (E.E.G.) and G20220125 (J.W.), Owens Family Glaucoma Research Fund, NEI grant R01EY017841 (E.E.G.), P30EY06360 (Emory Vision Core), and Challenge Grant from Research to Prevent Blindness.

## Availability of Data and Materials

The data quantifying axonal regeneration in the BXD mouse strain set is available on GeneNetwork (genenetwork.org): number of axons at 0.5mm (BXD_27559), number of axons at 1mm (BXD_27560), the distance traveled by the 5 longest axons (BXD_27561) and the length of the single longest axon (BXD_27562). All vectors used in this study are available upon request.

## Ethics Approval

All procedures involving animals were approved by the Animal Care and Use Committee of Emory University and were in accordance with the ARVO Statement for the Use of Animals in Ophthalmic and Vision Research.

## Competing Interests

The authors declare that they have no competing interests.

---

**Supplemental Figure 1.**
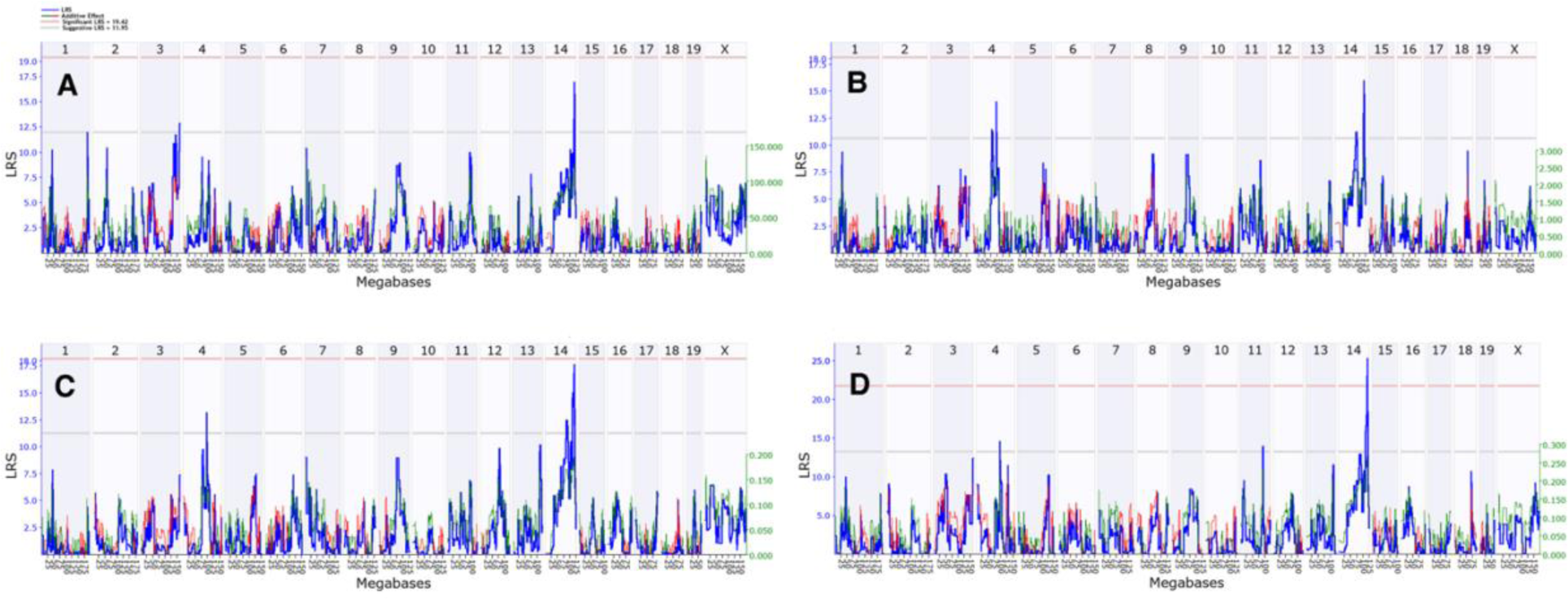
Genome-wide interval maps for the number of axons at 0.5mm from the crush site (A) and the number of axons 1mm from the crush site (B) are shown. In addition, the distance traveled by the longest 5 regenerating axons (C) and the longest single axon (D) are illustrated. The interval map plots the likelihood ratio statistic (LRS) across the genome from chromosome 1 to chromosome X. The light gray line is the suggestive level and the light red line is the genome-wide significance (*p=*0.05). When the optic nerve regeneration measures were mapped to the mouse genome, a significant association between regeneration and a locus on Chromosome 14 was observed.

## Notes

### Competing Interest Statement

The authors have declared no competing interest.

### Summary of Updates

We have changed the title to adequately define the nature of the manuscript. We have rewritten the abstract to more clearly define the relevance of the study. We have altered the introduction to make it clear to the reader the purpose and background of our manuscript. Minor editorial changes were also made in the manuscript.

